# Ephaptic coupling in white matter fibre bundles modulates axonal transmission delays

**DOI:** 10.1101/2020.04.08.031641

**Authors:** Helmut Schmidt, Gerald Hahn, Gustavo Deco, Thomas R. Knösche

## Abstract

Axonal connections are widely regarded as faithful transmitters of neuronal signals with fixed delays. The reasoning behind this is that extra-cellular potentials caused by spikes travelling along axons are too small to have an effect on other axons. Here we devise a computational framework that allows us to study the effect of extracellular potentials generated by spike volleys in axonal fibre bundles on axonal transmission delays. We demonstrate that, although the extracellular potentials generated by single spikes are of the order of microvolts, the collective extracellular potential generated by spike volleys can reach several millivolts. As a consequence, the resulting depolarisation of the axonal membranes increases the velocity of spikes, and therefore reduces axonal delays between brain areas. Driving a neural mass model with such spike volleys, we further demonstrate that only ephaptic coupling can explain the reduction of stimulus latencies with increased stimulus intensities, as observed in many psychological experiments.

**Author summary:** Axonal fibre bundles that connect distant cortical areas contain millions of densely packed axons. When synchronous spike volleys travel through such fibre bundles, the extracellular potential within the bundles is perturbed. We use computer simulations to examine the magnitude and shape of this perturbation, and demonstrate that it is sufficiently strong to affect axonal transmission speeds. Since most spikes within a spike volley are positioned in an area where the extracellular potential is negative (relative to a distant reference), the resulting depolarisation of the axonal membranes accelerates the spike volley on average. This finding is in contrast to previous studies of ephaptic coupling effects between axons, where ephaptic coupling was found to slow down spike propagation. Our finding has consequences for information transmission and synchronisation between cortical areas.

## Introduction

Signal processing and transmission in neuronal systems involves currents flowing across neuronal cell membranes. Due to the resistance of the extracellular medium, such transmembrane currents generate extracellular potentials (EPs), also called local field potentials (LFPs). The sources of EPs are synaptic currents, action potentials, calcium spikes and voltage-dependent intrinsic currents [1]. Neurons can therefore interact with their neighbours by changing the electric potential of the extracellular medium (and hence the membrane potential of their neighbours) without forming synapses. Such interaction is termed ephaptic interaction or ephaptic coupling [2–4]. Since EPs generated in the cortex are generally of the order of 100*μ*V [5] and therefore small in comparison to neuronal threshold potentials, the influence of EPs on neural computation is often regarded as negligible. EPs can be measured with intracranial electrodes and are used as a proxy for the underlying neuronal activity [6–9].

Seminal experiments by Katz and Schmitt [10], Rosenblueth [11], Arvanitaki [2] and Marrazzi and Lorente de Nó [12] have demonstrated that action potentials travelling along parallel axons can interact with each other if the extracellular medium is highly resistive. They demonstrated that action potentials with an initial offset would resynchronise, and also slow each other down. Furthermore, action potentials could be initiated in passive axons by action potentials travelling in a nearby axon. Several studies have reproduced these effects using computational models [13–20]. However, the experimental setup is such that the axons are placed into a highly resistive medium (either paraffin oil [10], or moist air [11]) in comparison to the intracellular medium, and the computational models assume that axons are embedded within a finite-sized extracellular medium. The latter would be justified by the presence of epineuria or perineuria, which is tissue restricting the extracellular space around axons in the peripheral nervous system. Both, however, are unlikely scenarios for axonal fibre bundles within the brain: the extracellular medium is only about three times more resistive than the intracellular medium, and axons in the CNS are not wrapped by epineuria and perineuria that would justify the ‘cables within a cable’ approach. For these reasons, the amplitude of extracellular potentials around spike carrying axons should be small, and ephaptic coupling should not play a significant role between individual pairs of axons within axonal fibre bundles in the brain. However, we hypothesise that collective interaction between multiple axons affects axonal signal transmission.

We test our hypothesis by introducing a modelling framework in which EPs modulate spike thresholds, and hence spike propagation velocities. We first determine the EPs generated by action potentials in single axons, which can be computed using the axial profile of an action potential (Fig 1a). The importance of computing the EPs lies in the fact that they perturb the membrane potential of a passive fibre (Fig 1b). This is then followed by the computation of EPs generated by spike volleys in fibre bundles (Fig 1c&d). As axon bundles contain millions of axons, we compute the cumulative effect of spike volleys at the macroscopic scale in axon bundles with diameters of several millimeters. The results of this analysis are used to build a point model for spike propagation, in which each spike travels with a velocity that is determined by structural parameters of the axon and the extracellular potential. This model is then coupled into a neural mass model to investigate *in-silico* the latency-intensity relationship of sensory stimuli, and the role of ephaptic coupling.

**Fig 1.**
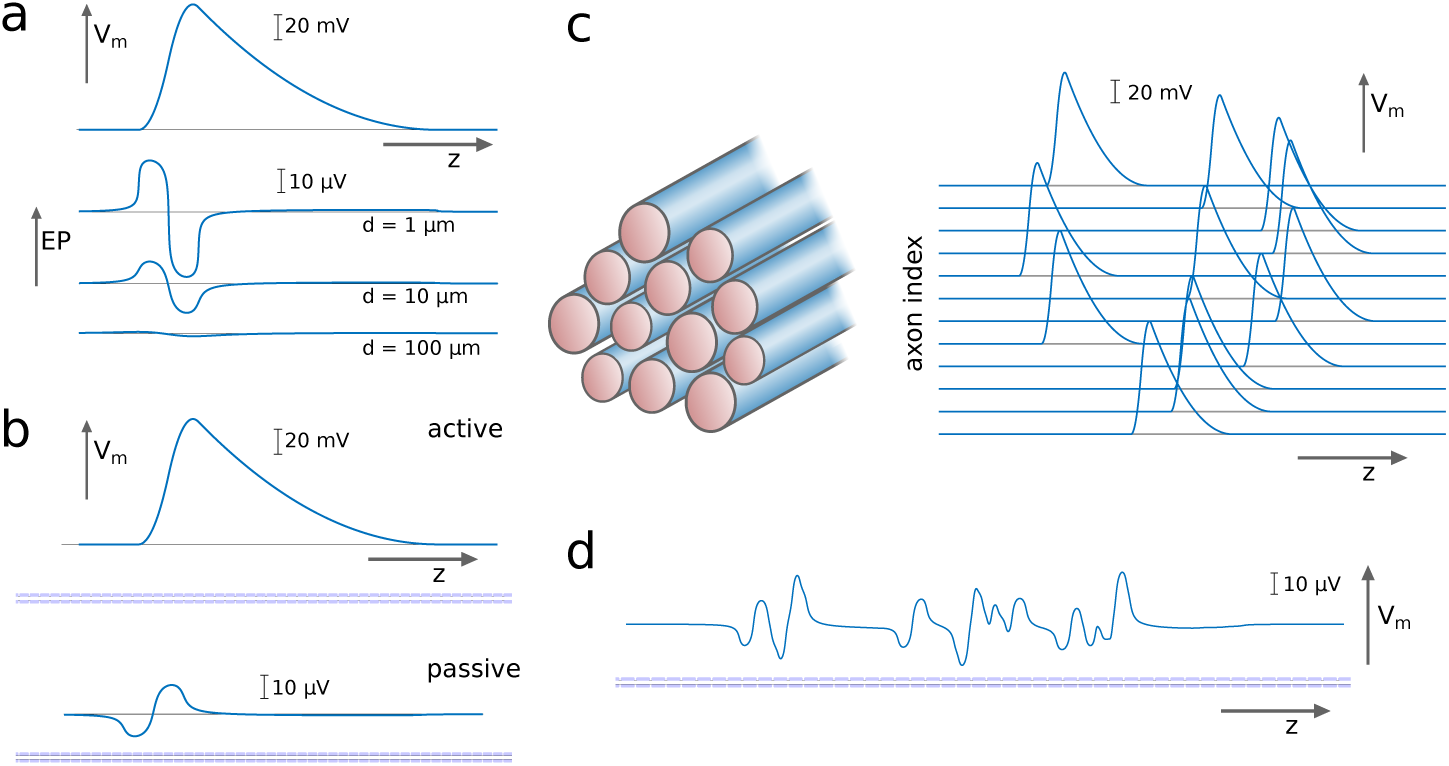
Computing the extracellular potential (EP) generated by a volley of spikes. (a) An action potential, as expressed by the membrane potential *V*_*m*_ along the axial dimension *z*, generates an EP that varies with *z* and the distance from the axon *d*. (b) An action potential in an active axon perturbs the membrane potential of a passive axon via the EP. (c) We consider spike volleys travelling along axonal fibre bundles, and (d) infer from the EP the cumulative effect on the membrane potential of a passive axon.

## Results

### Extracellular potentials around single axons

First we computed the EPs generated by action potentials in single axons. We use the line approximation [21–23], given that the diameters of axons are several orders of magnitude smaller than the diameter (or the general lateral dimensions) of axonal fibre

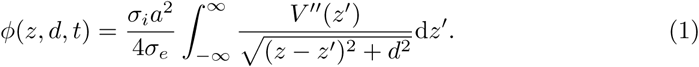

Here, *ϕ* is the EP, *z* is the axial dimension, *d* the distance from the axon, *t* is time, *σ*_*i*_ and *σ*_*e*_ are the intracellular and extracellular conductivities, *a* is the axon radius and *V* ‣(*z*) is the second derivative (curvature) of the membrane potential *V*. The EP is computed for different spatial profiles of action potentials, which include a piecewise linear and a piecewise quadratic approximation of spike profiles, but also spike profiles generated by a biophysical model [24] (Fig 2 (a-c)). The advantage of the piecewise approximation of the action potential profile is that the EP can be computed analytically (see the Methods section for details). The EP obtained from the biophysical model is computed numerically. For all the profiles we find that the maximum amplitude of the EPs is of the order of microvolts (Fig 2 (d-f)), and at long distances *d* the EPs decays with *d*^−3^ (Fig 2 (g-i)), akin to electric potentials of quadrupoles.

**Fig 2.**
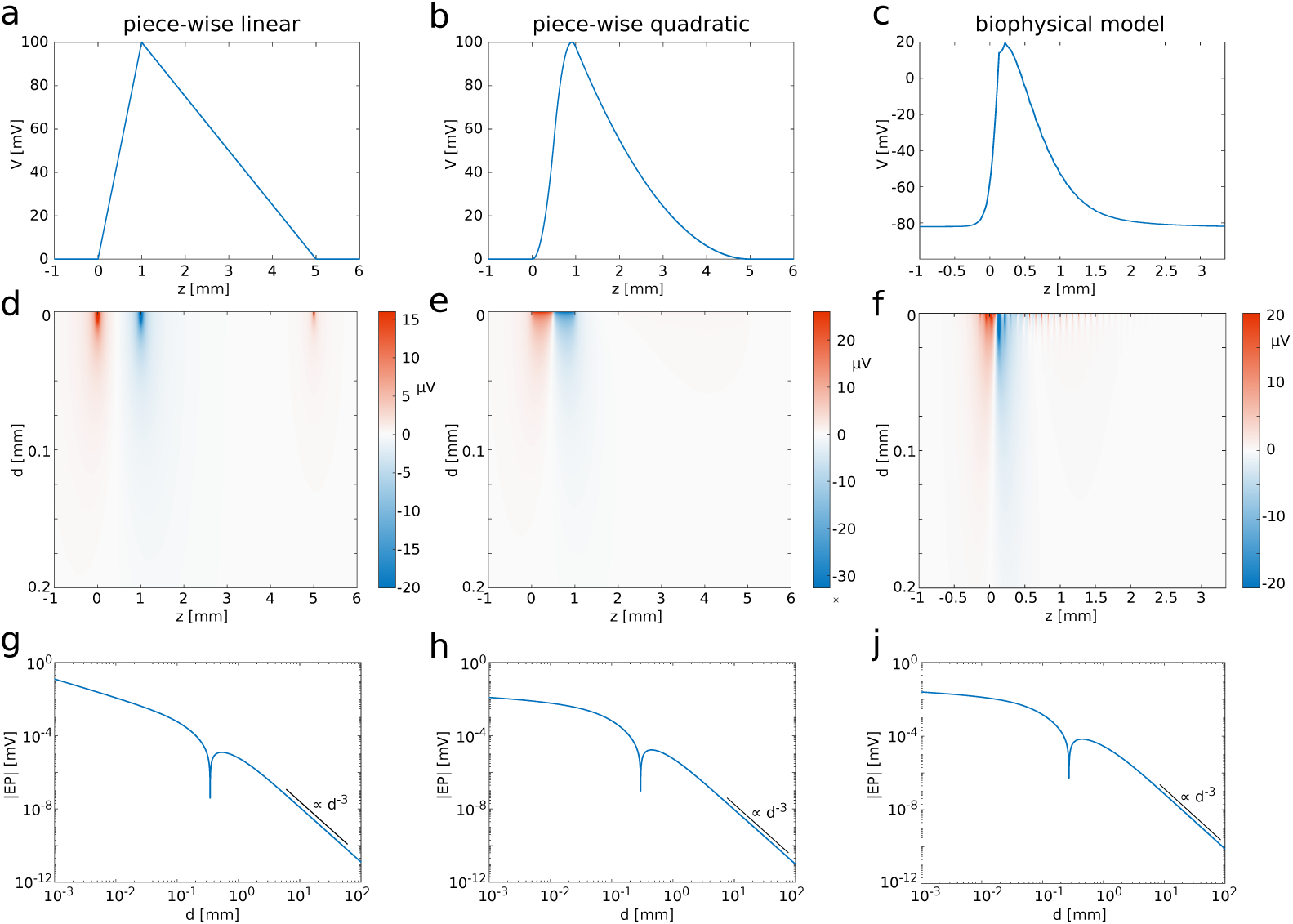
Spatial profiles of action potentials and their EPs. Shown are (a) the piecewise linear profile, (b) the piecewise quadratic profile, and (c) the profile of an action potential generated with the biophysical model. (d-f) EPs corresponding to action potential profiles in (a-c). (g-i) Log-log plots of the EPs (absolute values) at *z* = 0. Black lines indicate decay with *d*^−3^. (The notch at *d* ≈ 0.3mm is due to a change of sign.)

### Extracellular potentials in fibre bundles

To compute the effect of multiple action potentials in a fibre bundle, we assumed that a completely synchronous spike volley travels through the fibre bundle. The fibre bundle was arranged as a set of concentric rings of axons, as shown in Fig 3a. The reference point to compute the EP was set at the centre of the axon bundle. We computed the EP for an increasing number of spikes, beginning with six spikes in the innermost ring of axons, then 18 spikes in the two innermost rings, and successively increasing the number of rings in which all axons carry action potentials (Fig 3a). The maximum number of rings considered in this setup was 10^4^, which corresponds to a fibre diameter of 10mm if the diameter of the uniform axons is 0.5*μ*m. This fibre bundle contains approximately 3 × 10^8^ axons, similar to the number of axons in the corpus callosum [25].

**Fig 3.**
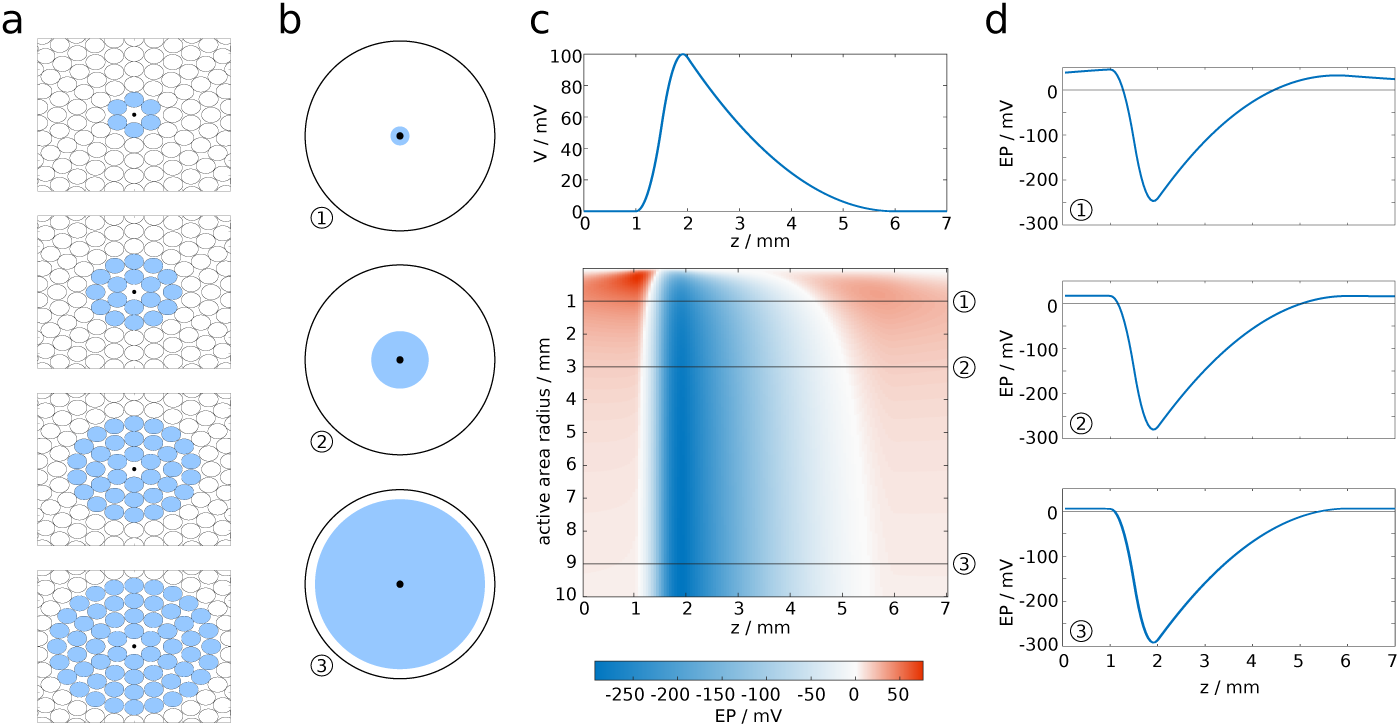
EP at the centre of a circular axon bundle due to concentric spike volleys. (a) Microscopic cross-section of a fibre bundle, with spike-carrying axons marked in blue. (b) Macroscopic extension of (a), with the active area (i.e. where axons carry spikes) marked in blue. (c) Spatial (longitudinal) profile of the EP at the centre of the fibre bundle. (d) Cross-sections of (c).

Increasing the active area (see Fig 3b for a macroscopic representation) yielded a longitudinal profile of the EP that saturated at large diameters (Fig 3c). Interestingly, the profile is approximately proportional to −*V*(*z*), with *V*(*z*) being the spatial profile of the action potential (Fig 3d). In the Methods section we demonstrate that this profile can be computed analytically, to a very good approximation, by the following expression:

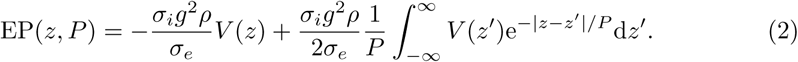

Here, *g* represents the ratio between the axon diameter and fibre diameter (axon plus myelin), commonly referred to as g-ratio. The relative size of the volume occupied by fibres, the fibre volume fraction, is represented by the quantity *ρ*, and *P* is the radius of the fibre bundle.

Next, we investigated how the EP changes with the position of the reference point, i.e. the point in the cross-sectional plane at which the EP is computed (Fig 4a). We found that the amplitude and longitudinal profile remained nearly unchanged, even if the reference point is close to the surface, as shown in Fig 4b&c. More specifically, the decrease of the amplitude is less than ten percent when the reference point is moved from the centre of the fibre bundle to 0.8 bundle radii away from the centre. Closer to the surface, the drop in amplitude is more marked. Outside of the bundle, while moving the reference point further away from the centre the EP drops rapidly, and at sufficiently large distances the drop in amplitude is proportional to *d*^−3^. We take this as evidence that the EP at the centre of the bundle is characteristic for the EP across the entire cross-section of the fibre bundle, i.e. we assume the EP is uniform in the radial direction.

**Fig 4.**
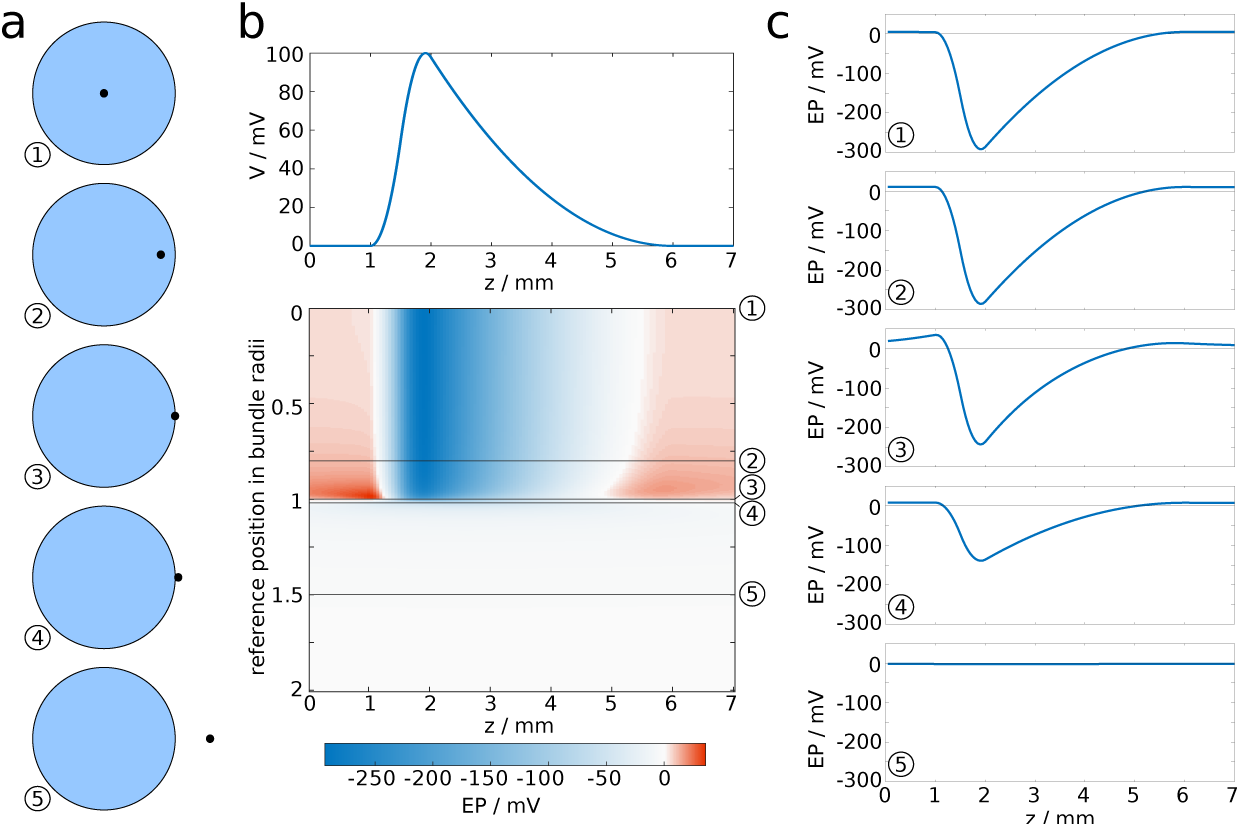
EP in fibre bundle with synchronous spike volley, subject to position of reference point. (a) The reference point is moved from the centre of the fibre bundle to a position outside of the fibre bundle. (b) EP plotted against the longitudinal coordinate *z* and the distance of the reference point from the centre. (c) Cross-sections of (b).

We consider spike volleys that engage all axons in the fibre bundle, which leads to EPs with amplitudes of order 100mV, as can be seen in Fig 3c and Fig 4b. This is certainly an unphysiological scenario, since it is unlikely that all axons in a fibre bundle carry perfectly synchronised action potentials, and because such large EPs would certainly disrupt signal transmission in the participating axons. However, it is plausible that a (sufficiently synchronous) spike volley might engage one percent of the axons in the fibre bundle, in which case the amplitude of the EP would be of the order of 1mV.

Alternatively, one may consider a spike volley that is not perfectly synchronised, i.e. the spikes are distributed in space due to varying emission times. To illustrate the effect of such a spatial distribution, we draw spike positions randomly from a uniform distribution of varying width Δ*z*. This spatial distribution can be associated with a temporal distribution via the relation Δ*z* = *v*Δ*t*, where *v* is the (intrinsic) propagation velocity of the uniform axons. In Fig 5 we show how increasing the active area affects the EP for different Δ*z*. It can be seen that the maximum amplitude decreases with increasing Δ*z*, and for sufficiently wide spike volleys the largest amplitude of the EP occurs near the edges of the spike volley instead of its centre.

**Fig 5.**
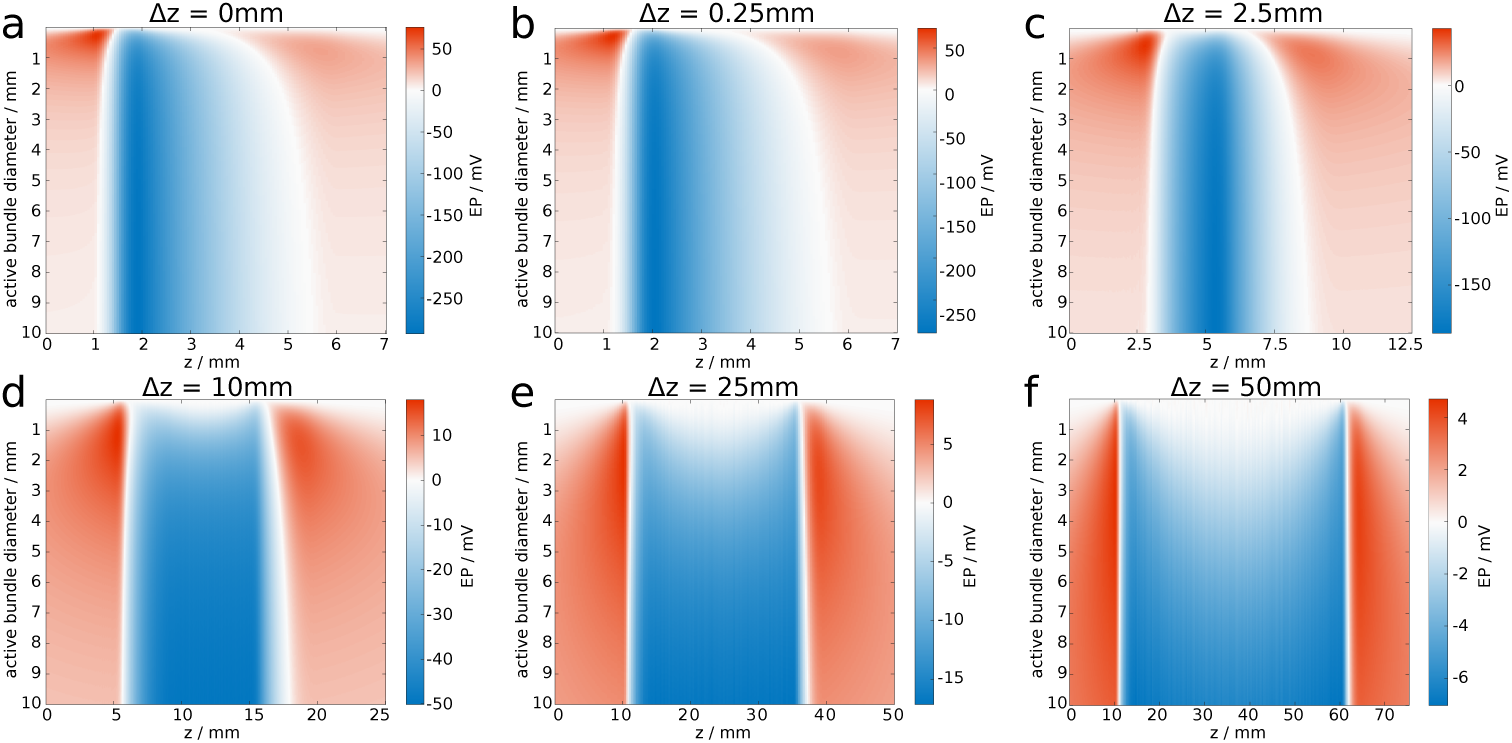
Increasing length of spike volley attenuates EP. The EP is shown for varying bundle diameters and *z*. We steadily increase the width of the spike volley from (a) to (f).

### A model for spike propagation

In addition to studying EPs generated by spike volleys in axonal fibre bundles, we are interested in the effect that EPs have on axonal signal transmission. Since the membrane potential is measured as the difference between intracellular and extracellular potential, a change of the EP implies a change of the membrane potential. For example, if the EP decreases, then the membrane potential increases, i.e. the membrane is depolarised. We assume that the EPs are not compensated by transmembrane currents or ionic shielding, or if such processes occur, that these processes are too slow to be relevant for short spike volleys.

We begin the modelling procedure by setting up a fibre bundle with *N* axons, each of which has a diameter drawn from a shifted alpha distribution that was chosen to closely fit the results by Liewald et al. [26] (Fig 6a). For numerical purposes we set the number of axons *N* between 10^3^ and 10^4^. A realistic fibre bundle contains many more axons, likely by several orders of magnitude. Conceptually, each of our model axons therefore represents a large number of axons with identical properties, but evenly distributed across the cross-sectional area. The fibre bundle is also endowed with macroscopic properties, namely the length and radius of the fibre bundle.

**Fig 6.**
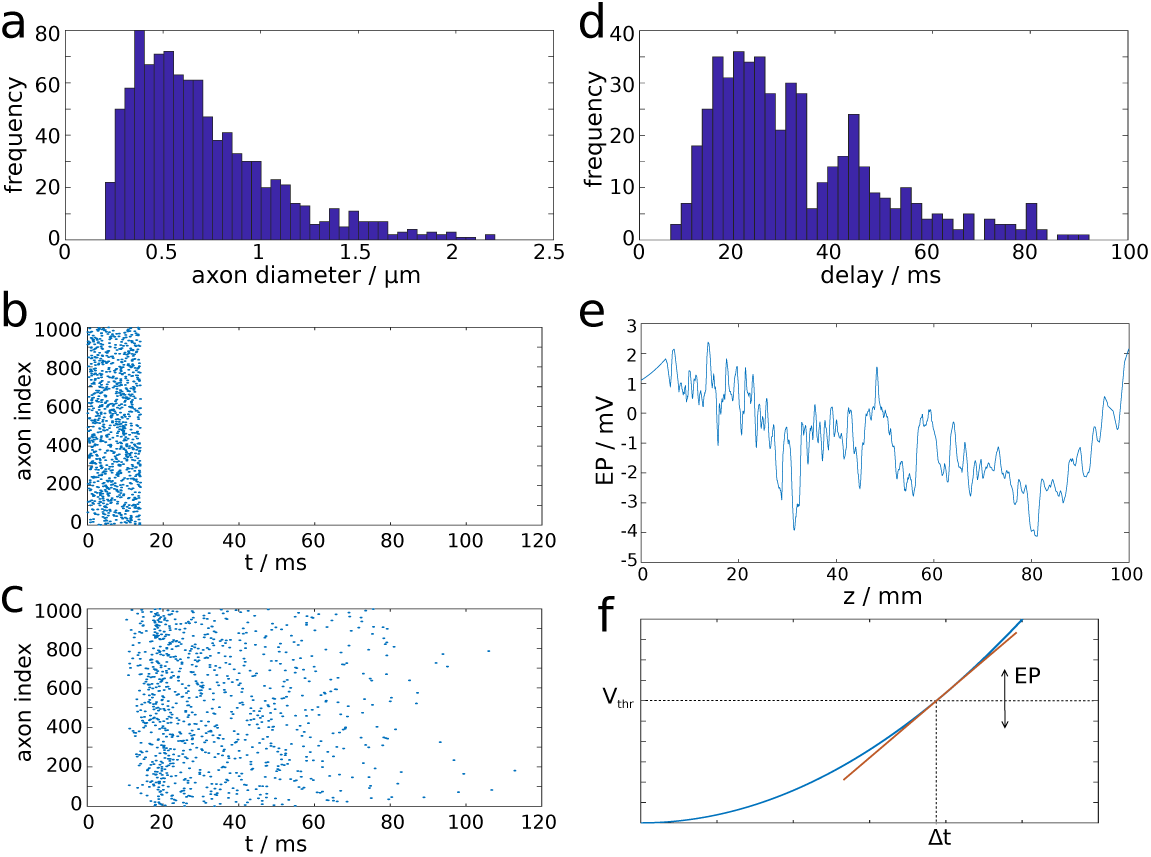
Illustration of properties of the computational model. (a) Distribution of axon diameters sampled from a shifted alpha distribution to match experimental data [26]. (b) Rastergram of spike volley generated at proximal end of fibre bundle. (c) Rastergram of spike volley reaching the distal end of the fibre bundle. (d) Distribution of delay times. (e) Longitudinal profile of EP generated by a spike volley. (f) The EP modulates the spiking threshold (*V*_*thr*_) and therefore the delay Δ*t* of action potential generation between two reference points (e.g. two consecutive nodes of Ranvier).

To test the transmission properties of a fibre bundle, we set up a spike volley with spike times drawn from a uniform distribution. The spike times define when the action potentials are generated at the proximal end of the bundle (Fig 6b). The propagation of spikes along the axon is determined by a point model that is described in the next paragraph. The spike volley then reaches the distal end of the fibre bundle (Fig 6c).

Due to the distribution of axon diameters, this process results in a distribution of transmission delays (Fig 6d). If the position of a spike is known, one can determine the EP generated by this spike. Since each model axon represents a large number of biological axons, we do not use the expression for single axons (Eq (1), but the one for the cumulative EPs generated by spike volleys (Eq (2)). The EP generated by a spike is thus the EP shown in Fig 3c, divided by the volume fraction occupied by the model axon. In this way, one can compute the spatial profile of the EP generated by a spike volley, see Fig 6e.

We employ a point model to track the position of an action potential along the fibre bundle. The position is determined by the leading edge (rising phase) of the action potential. For the linear and quadratic approximations of the spike profile, the position is defined by the point where the membrane potential first deviates from resting potential. In the absence of perturbations by non-zero EPs, the velocity is constant along the fibre bundle. Therefore, the position of a spike can be tracked by multiplying the intrinsic velocity (determined by structural parameters of the axon) with the time elapsed since the spike was generated. The velocity of a spike is also determined by a putative spike threshold, which might be interpreted within a spike-diffuse-spike framework [19]. It has been demonstrated, using some simplifying assumptions, that the spike threshold *V*_*thr*_ can be related to the activation delay Δ*t* between two subsequent nodes of Ranvier by some nonlinear function, and therefore to the velocity of a spike [19]. In the presence of EPs, the spike threshold, and therefore the propagation velocity, is perturbed. For simplicity, we assume a linear relationship between *V*_*thr*_ and Δ*t* (Fig 6f). This results in the following relationship between the perturbed propagation velocity *v* and the EP:

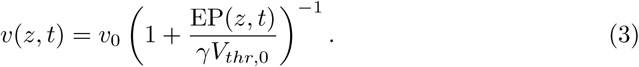

### Effect of extracellular potentials on transmission delays

The point model allows us to test the consequences of ephaptic coupling via EPs in a macroscopic fibre bundle. We investigate the dynamics of spike volleys with and without ephaptic coupling, and the resulting differences in axonal delays. There are several structural parameters that we keep fixed for simplicity, such as the fibre volume fraction (80% [27]), the fibre length (10cm), and the distribution of axon diameters. The spikes are generated at the proximal end of the fibre bundle, with spike times drawn from a uniform distribution. The width of this distribution determines the duration of a stimulus, and the number of spikes determines its intensity. We record the delays for each spike as it travels from proximal to distal end of the fibre bundle.

We first investigated how axonal delays are affected by ephaptic coupling, and focused on the mean of the delay distribution. In the presence of ephaptic coupling, we observe a decrease of the mean axonal delays as the stimulus intensity is increased (solid lines in Fig 7). In the absence of such coupling, the mean axonal delays remain constant (dashed lines in Fig 7). The stimulus duration is set to either 10ms or 20ms, and the bundle diameters are varied between 2mm and 8mm. The mean axonal delays drop nonlinearly with increasing intensity in the presence of ephaptic coupling, but remain unchanged in its absence (Fig 7 a&b). At full intensity (100%) and with ephaptic coupling, the mean axonal delays drop from 34ms to 24ms as the diameter of the fibre bundle is increased from 2mm to 8mm if the stimulus duration is 10ms (Fig 7a). At 20ms stimulus duration, the mean axonal delays drop from 36ms (unchanged) to 25ms with increasing diameter of the fibre bundle (Fig 7b). In other words, the mean axonal delay decreased by up to 30% at full stimulus intensity.

**Fig 7.**
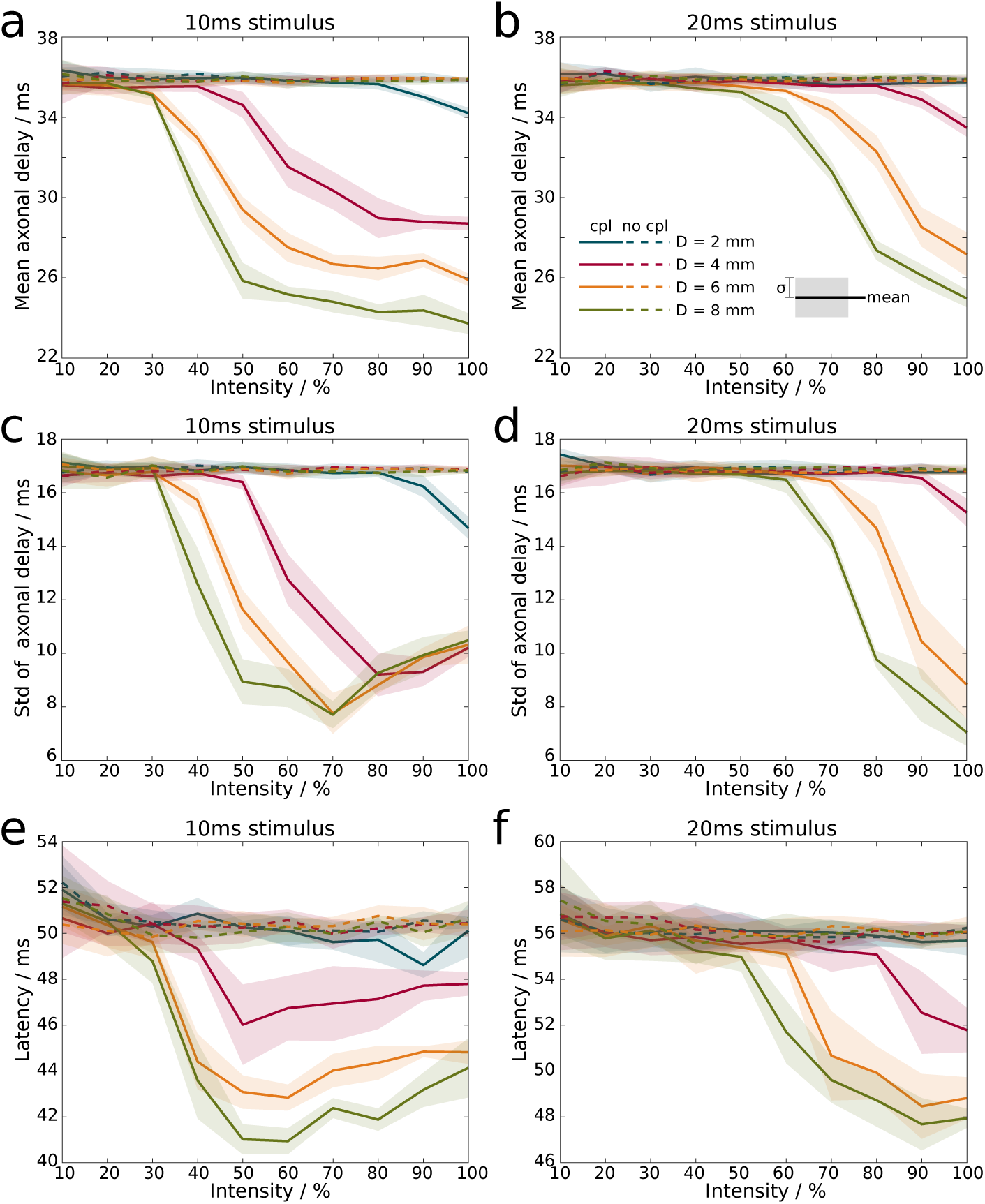
Increasing the stimulus intensity, i.e. the number of spikes in a volley, decreases axonal transmission times and the latency of stimulus response. (a) Mean axonal delay with ephaptic coupling (solid) and without ephaptic coupling (dashed) for 10ms stimulus duration, and (b) for 20ms stimulus duration. (c) Standard deviation from the mean of axonal delay with ephaptic coupling (solid) and without ephaptic coupling (dashed) for 10ms and (d) 20ms stimulus duration. Mean and standard deviation are computed from the distribution of delay times (cf. Fig 6d). (e) Latency from stimulus onset to first maximum in neural mass model at 10ms stimulus duration, and (f) at 20ms stimulus duration. Lines (shaded areas) indicate mean (1*σ* confidence interval) across 5 simulations. Colours indicate different bundle diameters.

Next, we explored how the standard deviation of axonal delays (a measure for its dispersion) behaved in the presence of ephaptic coupling. We found that its qualitative behaviour is similar to the mean of the axonal delays (Fig 7 c&d), with a nonlinear decrease as the stimulus intensity is increased. However, the relative decrease of the standard deviation is even more marked than the decrease of the mean, which indicates that ephaptic coupling reduces dispersive effects (that occur, for example, due to different fibre diameters), resulting in more synchronous spike volleys than in the absence of ephaptic coupling.

Finally, we incorporated the axon bundle into the Jansen-Rit neural mass model [28]. The arrival of each spike at the distal end generates a current that is injected into the neural mass model. The response latency is determined by the time difference between stimulus onset and the maximum response of the neural mass model. This results in increased latencies as the stimulus duration is increased. However, in the presence of ephaptic coupling, we observe again a nonlinear decrease in latencies as the stimulus intensity in increased, whereas in the absence of ephaptic coupling the decrease is only marginal (Fig 7 e&f). Regardless of stimulus duration, at full stimulus intensity ephaptic coupling reduces the response latency by up to 8ms, which corresponds to a reduction by approximately 15%.

## Discussion

The key finding of our study is that spike volleys generate EPs with sufficiently large amplitudes to modulate axonal delays. Specifically, the mean delay of a spike volley decreases as the number of spikes in the spike volley is increased. Therefore, our results suggest that varying the amplitude of a neuronal signal can adjust its delay. Using a neural mass model, we have demonstrated that the decrease of axonal delays translates into the decrease of stimulus latencies as the stimulus intensity is increased. Our modelling approach required several assumptions that we are going to discuss in more detail.

We computed the EPs using the line approximation (i.e. the axon is assumed to be infinitely thin), which has been demonstrated to be very accurate [21]. We further assumed that the axon bundle is large, circular, homogeneous, and densely populated with axons. The latter is justified by electron micrography studies which suggest that only a small fraction of an axon bundle is made up of extracellular space [26, 29]. Since axonal membranes and the myelin sheaths have a much larger resistivity compared to the extracellular medium, electric currents can only pass through the extracellular medium. We assumed that the medium is homogeneous and that the effective conductivity of the fibre bundle is the conductivity of the extracellular fluid multiplied by the relative size of the extracellular space. This calls for more detailed simulations of the spread of EPs with spatial heterogeneity taken into account. For mathematical convenience, we chose the fibre bundles to be large with circular cross sections. Realistic fibre bundles are indeed large, but often show a more sheet-like morphology [30]. An open question is whether this morphology influences the effect of EPs within our framework (a recent study used coupled axons with FitzHugh-Nagumo dynamics to demonstrate ephaptic coupling effects in sheet-like bundles [20]).

Furthermore, we ignored possible effects due to the axonal microstructure. We assumed the axonal membrane to be smooth (effectively a homogenised axon [31]), and that therefore nodes of Ranvier are not relevant as point sources. This is certainly the case at large distances from the axon, as can be seen in Fig 2f. However, at close proximity such effects would be relevant, as the EP at a node of Ranvier can reach several hundreds of microvolts. It is unknown whether nodes of neighbouring axons are sufficiently aligned to affect action potential generation in such a manner. As oligodendrocytes can myelinate multiple axons [32, 33], it is conceivable that neighbouring axons show some degree of alignments, in which case it would be possible to observe ephaptic coupling effects in much smaller fibre bundles, provided the spike volleys are sufficiently synchronised.

To demonstrate the effect of EPs on axonal delays we used a strongly simplified model for spike propagation. This model assumes that the spike velocity resulting from the axon morphology is known, and that this velocity is perturbed by the EP. It does not contain possible compensation effects arising from Hodgkin-Huxley dynamics (i.e. subthreshold currents that repolarise the axonal membrane), and further studies are required using the Hodgkin-Huxley framework to confirm our results. We note that such an undertaking would be computationally extremely expensive, and is therefore beyond the scope of the present study.

We have incorporated the axon bundle model into the Jansen-Rit neural mass model to build a model system for primary sensory information processing, and to investigate the relationship between stimulus intensity and response latency. Psychological experiments across different sensory modalities yield the same qualitative relationship, whereby the latency decreases with increasing intensity [34–40]. Such experiments typically measure the delay between stimulus presentation and the first maximum or minimum of the neural response measured electrographically. To replicate this experimental design, we measured the time difference between stimulus onset, i.e. the start of the spike volley, to the first maximum in the response of the Jansen-Rit model. Interestingly, only the presence of ephaptic coupling could explain the latency-intensity relationship. We are aware that the Jansen-Rit model is a fairly simple representation of a cortical microcircuit, and that other nonlinear processes not taken into account in its derivation may also reduce the response latency with increased stimulus intensity, such as oscillation-mediated information transmission [41]. Nevertheless, our modelling approach suggests that ephaptic coupling effects play a role in neural responses.

While there is such implicit evidence, further experimental studies are necessary to test our hypothesis. The experimental design would be highly invasive, since the EPs drop rapidly with distance outside fibre bundles. Animal experiments have already demonstrated the possibility to record EPs within axonal fibre bundles [23, 42]. An interesting test bed could also be a delay analysis within stimulation-response paradigms used in epileptic patients to determine the seizure focus [43, 44].

If such activity-dependent (or rate-dependent) delays occur in fibre bundles, then one may speculate as to their putative role in information processing. Since axonal delays are in general quite small (about 30ms in a 10cm long fibre bundle), the main effect should be on fast oscillations. It is indeed tempting to propose that such variable delays may have an effect on long-range gamma synchronisation, and that synchronisation patterns can be flexibly switched by changes in the amplitude of the transmitted spike volleys. We have found that ephaptic coupling can decrease delays by up to 10ms, which would be one half of the period of a gamma cycle at 50Hz. It has been demonstrated that delays are critical in shaping the functional architecture of the brain [45–47], and ephaptic modulation of such delays could therefore play a role in flexibly synchronising distant brain areas.

## Methods

In this section, we describe the mathematical framework underlying our study. We use a detailed microscopic description of the interaction between axonal fibres, and a leading-edge approximation which reduces the computational effort, but retains key properties of the interaction. We investigate a fibre bundle in which axons are coupled by the EPs generated by spikes. We first show how to compute the EPs generated by single spikes and spike volleys, and then present the framework of the point model for spike propagation.

### Single fibre

In an open fibre bundle, the EP is determined by currents entering and leaving an axon. The axial currents around an axon can be inferred from the spatial profile of an action potential [23]:

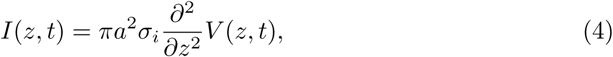

with *I*(*z, t*) being the axial (intra-axonal) currents, *V* (*z, t*) the membrane potential, *a* the axon radius and *σ*_*i*_ the conductivity of the intracellular medium. The axial dimension is represented by *z*, and time by *t*. The EP, denoted by *ϕ* for the single spike, can then be computed from the axial current via:

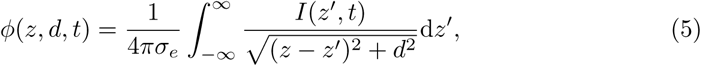

with *σ*_*e*_ being the extracellular conductivity, and *d* the radial coordinate measuring the distance from the axon. Inserting Eq (4) into Eq (5) yields

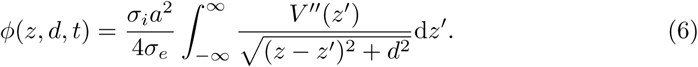

This integral is the convolution of the curvature of the action potential profile with the kernel *G*(*z* − *z*′) = (*z* − *z*′)^2^ + *d*^2^. In general, this integral has to be evaluated numerically. In order to obtain an analytical solution, we approximate the shape of an action potential by piecewise linear or piecewise quadratic functions.

#### Piecewise approximation of action potential profile

In general the profile of an action potential has to be determined either numerically, or using spike-diffuse-spike formalisms. In the former case it is impossible to parameterise the profile, and in the latter the analytical expressions are still prohibitive to follow through with the calculations of the EP. Therefore, we present a formalism which approximates the profile of an action potential with either piecewise linear or piecewise quadratic functions. This method can be extended to arbitrary polynomial expressions, and is similar to curve-fitting with splines.

##### piecewise linear approximation

The simplest approximation of an action potential is given by two linear functions on two consecutive intervals, describing the rising and the falling phase of the action potential, respectively:

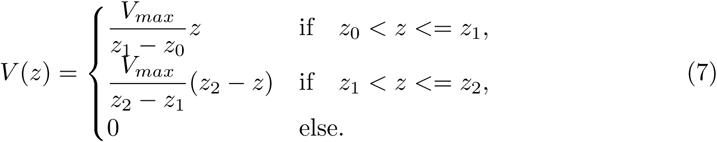

The first derivative of this approximation is piecewise constant with discontinuities at *z* = *z*_0_, *z* = *z*_1_ and *z* = *z*_2_. The second derivative is therefore

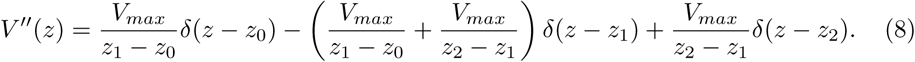

It is then straightforward to compute the EP:

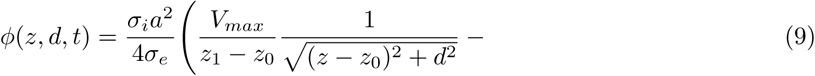

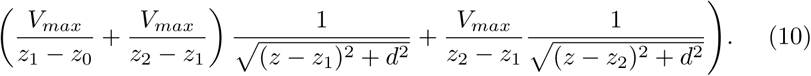

##### piecewise quadratic approximation

For the piecewise quadratic approximation, we divide the AP profile into three segments:

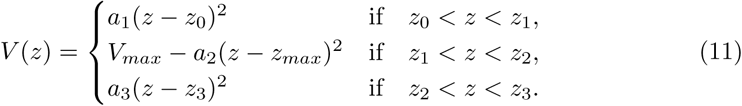

Given *z*_0_, *z*_1_, *z*_2_, *z*_3_ and *V*_*max*_, there are four unknowns *a*_1_, *a*_2_, *a*_3_ and *z*_*max*_. To ensure a smooth profile, we impose boundary conditions that assume *V* (*z*) is smoothly differentiable, i.e. 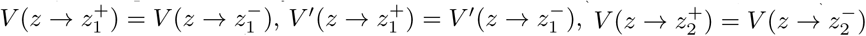, and 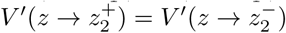. After some manipulation, we obtain:

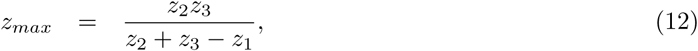

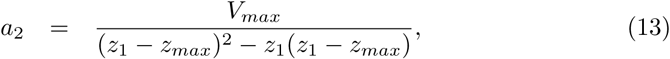

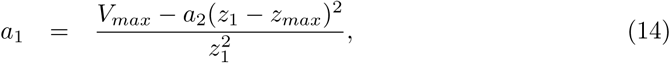

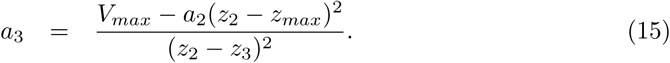

The second derivative of the spatial profile is piecewise constant:

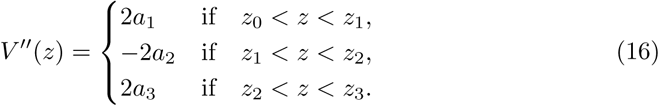

The EP is then found to be

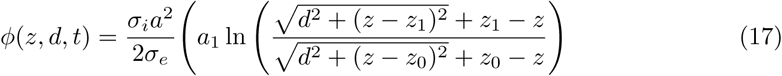

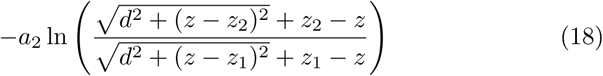

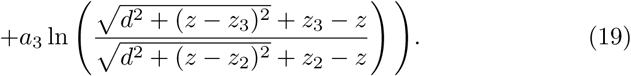

#### Biophysical model

In order to compare the piecewise approximations of spike profiles with the profile of a biophysically realistic spike, we used the model presented in [24] to obtain such a profile numerically. We used the code made available by the authors on github [48].

Specifically, we used the cortex model with standard parameters, and extracted the profile at a suitable time point when the membrane potential at both ends of the axon was close to resting potential. The EP was computed by solving Eq (1) numerically.

### Fibre bundle

#### bundle with identical axons

To compute an upper boundary of the EP produced by multiple action potentials, we assume that a perfectly synchronous spike-volley travels through a dense fibre bundle. All axons in this fibre bundle have the same diameter and are arranged in concentric rings (Fig 3a). At the centre of an empty grid position we compute the EP by summing *ϕ* at distance (2*n* + 1)*a* of 6*n* axons, with *n* ranging from 1 to *N*, with *N* large:

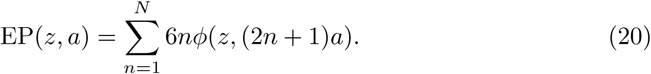

Although *ϕ* is approximately 20*μV* at the surface of an isolated axon, in a fibre bundle the combined effect can lead to EPs of many mV. Interestingly, we find that for large enough fibre bundle diameters the profile of the cumulative EP is almost proportional to the profile of the generating action potentials. We give a mathematical explanation for this next.

#### Analytical solution

The cumulative EP at the core of an axon bundle is computed with the following integral,

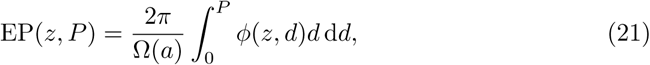

with *P* being the axon bundle diameter, and Ω(*a*) being the cross-sectional area occupied by an axon with radius *a*. We set Ω(*a*) = *πa*^2^/(*ρg*^2^), where *g* is the g-ratio and *ρ* is the fibre volume fraction.

Inserting Eq (1) into Eq (21), and solving the integral over *ρ*, results in

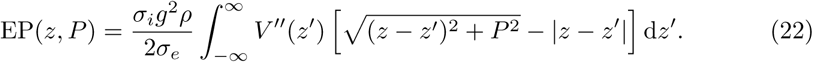

Integration by parts then yields

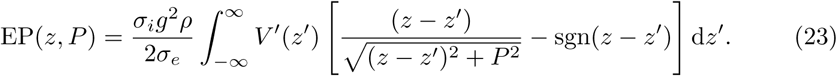

Next, we use the approximation

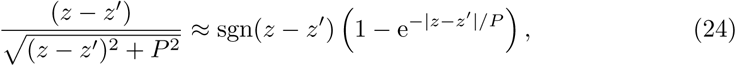

which leads to

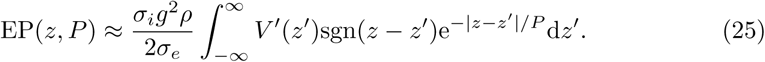

Using integration by parts of this integral ultimately yields

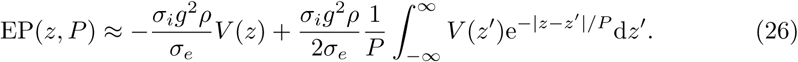

We may regard this result as the far-field approximation of EPs in axonal fibre bundles.

We note here also that in the limit *P* → 0, exp(|*z* − *z*′ |*/P*)*/P* → 2*δ*(*z* − *z*′), and yields

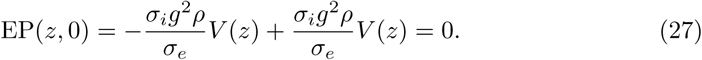

Eq (22) suggests that the far-field approximation of the cumulative EP is independent of the axon morphology. At this point, however, we have not taken into account that axons of different diameters transmit action potentials at different velocities, and that therefore the spatial profile widens with increasing action potential velocity.

### Leading edge model

The two most common ways to model axonal signal transmission are either Hodgkin-Huxley type dynamics embedded in a core-conductor model, or simpler spike-diffuse-spike approaches. Both ways allow one to determine the spike velocity as a function of electrophysiological and structural parameters. Here, we employ a much simpler model that describes the position *z*_*i*_ of an action potential (more precisely, its leading edge or rising phase) travelling along the *i*^*th*^ axon by one simple equation:

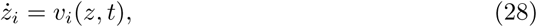

where *v*_*i*_(*z, t*) is the velocity of the action potential as function of the axial direction *z* and time *t*. If the axon is homogeneous and does not experience spatial or temporal perturbations, then the velocity can be expressed by *v*_*i*_(*z, t*) = *v*_*i*,0_, which is the intrinsic velocity of the axon, determined by its morphological and electrophysiological properties. We assume here that this velocity is known for each axon. In the absence of perturbations, one can therefore express the axonal delays by *τ*_*i*_ = *L/v*_*i*,0_, with *L* being the length of the fibre bundle. We set here *v*_*i*,0_ = *αd*_*i*_, with *d*_*i*_ being the diameter of the *i*^*th*^ axon, and *α* = 5ms^−1^*/μ*m.

Changes in the EP lead to perturbations of the membrane potential of an axon. A negative (positive) EP effectively depolarises (hyperpolarises) the axonal membrane, and therefore increases (decreases) the propagation velocity. A convenient formalism to incorporate such changes is the spike-diffuse-spike framework, in which the spiking threshold is a parameter to explicitly describe the onset of an action potential [19]. Such thresholds can also be determined within the Hodgkin-Huxley formalism, albeit these thresholds vary with the depolarisation rate [49]. The EP can therefore be regarded as a perturbation of such a threshold. Within the spike-diffuse-spike framework, the following relationship between the spike threshold *V*_*thr*_ and the delay of spike generation Δ*t* between two consecutive nodes of Ranvier can be derived:

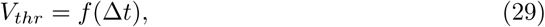

see Fig 6f for a visual representation. The function *f* (Δ*t*) depends and structural and electrophysiological parameters of the axon. The EP can be incorporated into the spiking threshold, *V*_*thr*_(*z, t*) = *V*_*thr*,0_ + EP(*z, t*), with *V*_*thr*,0_ being the uniform spiking threshold of the unperturbed axon. Via Eq (29) one can relate *V*_*thr*,0_ to Δ*t*_0_ of the unperturbed axon, and to its intrinsic velocity via *v*_*i*,0_ = *l/*Δ*t*_0_, where *l* is the distance between two consecutive nodes of Ranvier. To further simplify our scheme we linearise Eq (29) around *V*_*thr*,0_ and Δ*t*_0_:

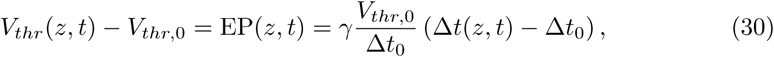

which can be reformulated into

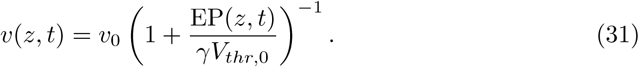

Here we made use of *v* = Δ*z/*Δ*t*(*z, t*), which results in Δ*t*(*z, t*)*/*Δ*t*_0_ = *v*_0_*/v*(*z, t*). The parameter *γ* denotes the relative steepness of *f* (Δ*t*) around Δ*t*_0_, and for typical threshold values can be set to *γ* = 2 (compare Fig 11 in [19]). This parameter can be used as a tuning parameter for the inverse strength of the ephaptic coupling, and in the numerical simulations we set *γ* = 10^3^ to represent the absence of any ephaptic interaction.

The variable *v*(*z, t*) represents the velocity of the leading edge of the spike. In order to compute the EP generated by an action potential, we need to know the length of the entire action potential. A convenient way is to relate the length of an action potential to its effective velocity, i.e. the velocity of its centre of mass. This effective velocity is, in essence, a time-averaged quantity. We approximate the effective velocity of a spike by

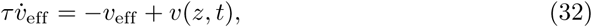

with *τ* = 1ms.

### Jansen-Rit microcircuit

The spike volleys represent cortical, subcortical or sensory information being transitted by axon bundles. To describe the response of neuronal circuits, e.g. cortical microcircuits, we use the Jansen-Rit model [28, 50, 51] and record the maximum response in the membrane potential of its pyramidal cell population. The Jansen-Rit model is composed of six differential equations:

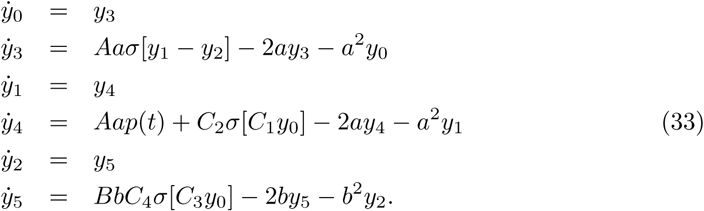

Here, *y*_0_ is the postsynaptic potential (PSP) generated by the output of the pyramidal cells at the two interneuron types, and *y*_1_ and *y*_2_ are the excitatory and inhibitory PSPs generated at the pyramidal cells by external stimuli *p*(*t*) and the firing activity of the interneurons. Furthermore, *y*_3_, *y*_4_ and *y*_5_ are auxiliary variables in the synaptic conversion of firing rates into PSPs, with *a* and *b* being the inverse time constants of excitatory and inhibitory synapses, and *A* and *B* the respective maximum amplitudes of the synaptic response. The PSPs are converted into firing rates by the sigmoidal function

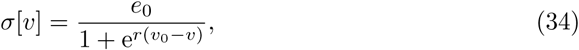

with *e*_0_ being the maximum firing rate, *v*_0_ the membrane potential at half of the maximum firing rate, and *r* sets the steepness of the sigmoid. The interaction between the different neuron types is scaled by the connectivity constants *C*_1_ to *C*_4_. The parameters are chosen as in [28]: *A* = 3.25mV, *B* = 22mV, *a* = 100*s*^−1^, *b* = 50*s*^−1^, *v*_0_ = 6mV, *e*_0_ = 5*s*^−1^, *r* = 0.56*mV* ^−1^, *C*_1_ = 135, *C*_2_ = 0.8*C*_1_, *C*_3_ = 0.25*C*_1_, and *C*_4_ = *C*_3_.

The external firing rate *p*(*t*) is generated by the incoming spikes,

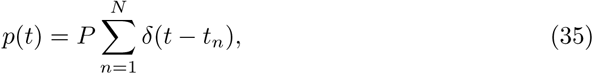

with *t*_*n*_ being the arrival time of the *n*^*th*^ spike, *N* the total number of spikes, and *P* = 0.1*s*^−1^ sets the coupling strength of the spike train.

The membrane potential *y* of the pyramidal cell population is determined by the difference between excitatory and inhibitory PSPs, i.e. *y* = *y*_1_ − *y*_2_. The response latency is then calculated as the position of the absolute maximum of *y*.

## Acknowledgments

HS, GH and TRK were supported by the German Research Foundation (DFG [KN 588/7-1] within priority programme ‘Computational Connectomics’ [SPP 2041]). GD was supported by the Spanish Research Project COBRAS PSI2016-75688-P (AEI/FEDER, EU), and by the Catalan AGAUR program 2017 SGR 1545.

